# Non-contact direct sensing of material properties of biomolecular condensate using Scanning Ion Conductance Microscopy

**DOI:** 10.64898/2026.06.16.731580

**Authors:** Helena Miljkovic, Karl Pang-Yeo, Zahra Ayar, Edoardo Fatti, Akhil S. Naidu, Jialin Shi, Marcos Penedo, Karsten Weis, Wayne Yang, Aleksandra Radenovic

## Abstract

Biomolecular condensates are important regulators of cellular compartmentalization and biochemical processes. Understanding their material properties is critical to elucidate how they control molecular organization and dynamics within cells. However, quantitatively probing these properties remains challenging due to the wide range of length scales, concentrations, and timescales over which condensates operate, as well as the limited force ranges accessible to current nanoscale mechanical mapping methods. We explored the use of a non-contact 3D imaging tool Scanning Ion Conductance Microscopy (SICM) for stiffness measurements of liquid–liquid phase-separated biomolecular condensates. We focus on the Dhh1 protein, which is a regulator of cytoplasmic processing bodies (PBs) membrane-less cytoplasmic condensates that control the storage and degradation of untranslated mRNA. In our study, we investigate the properties of mCherry2- or His-mCherry2-tagged full-length Dhh1 and N- or C-terminus tail-deletion constructs, as well as the catalytically inactive mutant DQAD, under different pH and incubation times. We mapped both spatial and temporal changes in the material properties of the condensates, highlighting the capabilities of the instrument. We found that the removal of either of the two tails led to an increase in condensate stiffness upon shifting the pH from a stress-associated cellular environment (pH 6.5) to physiological conditions (pH 7.5). Additionally, the choice of protein tags led to vastly different results depending on the pH where mCherry2-Dhh1 exhibited a stiffening going from pH 6.0 to 6.5 while the double-tagged His-mCherry2 did not. Our measurements are verified and corroborated by established techniques such as optical tweezer-based fusion assays and fluorescence recovery after photobleaching (FRAP). Furthermore, we were able to track the same biomolecular condensate sample for up to 20 days getting insights on the ‘ageing’ and evolution of the condensates. Overall, our study demonstrates the applicability of SICM for direct measurement of the material properties of biomolecular condensate.

## Introduction

Biomolecular condensates play a crucial role in cellular compartmentalization and the regulation of many biochemical processes^1,2^. The formation of biomolecular condensates is a process governed by local protein concentration, molecule interactions, as well as buffer conditions, notably pH and ionic strength, and the presence of other molecules such as RNA, DNA, etc. Within the condensates, proteins and polymers like DNA and RNA form interconnected, network-like structures where proteins multivalently interact with themselves or other proteins and biopolymers. This process is often reversible allowing for movement, reaction and subsequent transportation and release of the needed molecules post-processing. In recent years, biomolecular condensates have also been increasingly linked to the pathology of neurodegenerative diseases^3,4^, particularly when they undergo irreversible stiffening through a gel/solid-like transition^5,6^. Techniques to characterize material properties of biomolecular condensates are pivotal not only for their functional characterization but potentially also for diagnosis and therapy.

A plethora of techniques has been developed to probe the material properties of biomolecular condensates^7,8^. These techniques include optical tweezer fusion assays^7,9^, where the coalescence time between two condensates is measured (typically reported as τ-fusion time), and microrheology, which relies on tracking the trajectories of embedded beads to infer their viscoelastic properties^8^ or inducing them using AFM tips^10,11^ or optical tweezers^4,12–14^. There are also optical based techniques such as fluorescence recovery after photobleaching (FRAP)^15,16^, to recover diffusion coefficients and differential dynamic microscopy (DDM) to track intensity fluctuations in Fourier space through time. However, there are certain drawbacks to these techniques. Optical tweezers can introduce local heating and potential photodamage to proteins, while both optical tweezers and active micro-rheology typically suffer from low throughput because each condensate must be manipulated individually. With passive micro-rheology methods, both the size and the number of beads embedded in the condensate can influence the accessible measurement range and alter the measured properties. For FRAP and DDM techniques, the protein must be fluorescently labeled, which can lead to unwanted changes in condensation properties^17,18^. Additionally, most of these techniques give only qualitative measures of the material properties (e.g. the fusion time is a ratio of viscosity and surface tension), and the recovery time is a measure of the diffusivity rather than a direct quantity for stiffness^8^. Nevertheless, these techniques have served as qualitative indicators for changes in the material properties in numerous studies especially when comparing relative changes across different conditions.

In this work, we demonstrate a Scanning Ion Conductance Microscopy (SICM) based technique to directly obtain the stiffness of biomolecular condensates^19–21^. SICM is a non-contact topography imaging tool to map surfaces offering the advantage of ionic conductance flow rather than direct interaction with a tip such as in atomic force microscopy^22–28^. Additionally, this technique has recently been demonstrated to be able to measure stiffness and quantify the Young’s modulus by applying a pressure-induced flow to the sample and analyzing its mechanical response^20,19,21,29^.

In our case study, we employ SICM to directly probe the material properties of condensates composed of Dhh1 and RNA. Dhh1 is part of the family of DEAD-box ATPases^30^ and plays a crucial role in the dynamics of Processing Bodies (P-bodies), cytoplasmic condensates that function in cytoplasmic mRNA metabolism^31^. Structurally, Dhh1 is composed by a core domain composed by two RecA-like domains that mediate RNA binding and two unstructured tails, which are involved in condensate modulation. *In vivo*, Dhh1, together with untranslated mRNA and other proteins, i s required for P-body formation. In yeast, stresses such as acute glucose deprivation promote rapid P-body formation, whereas re-addition of glucose results in their fast dissolution^31^. Previous reports showed that *in vitro* condensation of Dhh1 with RNA is pH dependent, with lower pH favoring condensation and reducing condensate dynamics^17^. In this study, we investigate the material properties of full-length Dhh1 or variants devoid of the unstructured tails in different pH. Additionally, we explore the influence of commonly used mCh2 and His tag. Together, our study demonstrates the suitability of SICM to study the material properties of condensates and monitor their long timescale evolution over multiple days.

## Results

### Quantification of the stiffness by SICM

Typically, SICM is operated in an non-contact imaging tool where a potential difference is applied across the nanopipette, generating an ionic current that passes through the pore (i.e. the open nanopipette current)^23^. When the nanopipette is in proximity to a surface or boundary (∼ 600nm – 1µm^32,33^ depending of the size of the capillary and buffer), the ionic current is reduced due to obstruction of ionic flow. This reduction triggers the system and records the position of the nanopipette allowing for the reconstruction of the surface topography (height map of the sample)^32^. The ionic current moving through the nanopipette also induces a small electroosmotic flow of water toward the outside of the nanopipette, generating a local pressure on the sample surface. This local pressure can be changed either by physically bringing the probe closer to the sample or by changing the potential difference across the nanopipette thereby changing the electroosmotic flow^21^. The sample’s mechanical response (determined from the slight deformation of the surface) to the controlled pressure variation provides a means to quantify its local stiffness as it will in turn affect the change in the open nanopipette current as a function of height^20,21^. In practice, the nanopipette is brought gradually toward the surface using a combination of a stepper motor for coarse positioning and a piezoelectric actuator for fine control in the nanometer range. During this approach, the ionic current i s continuously monitored, and a predefined current threshold is used as a criterion to halt the motion, preventing contact and avoiding damage to the sample and the probe.

We performed SICM topography measurements on Dhh1 condensates prepared as described in the **Method section**. Briefly, Dhh1 was diluted to a concentration of 5 µM in a glass-bottom petri dish by the addition of the RNA substrate polyuridylic acid (pU) to trigger the formation of condensates (**Figure 1 A**). Condensates were allowed to settle to the bottom of the glass cover slip and were first inspected by transmitted light imaging through the inverted objective (**Figure 1B**). Optical images of the condensates were acquired before SICM measurements to document their morphology and position (see **Figure 1B**).

**Figure 1.**
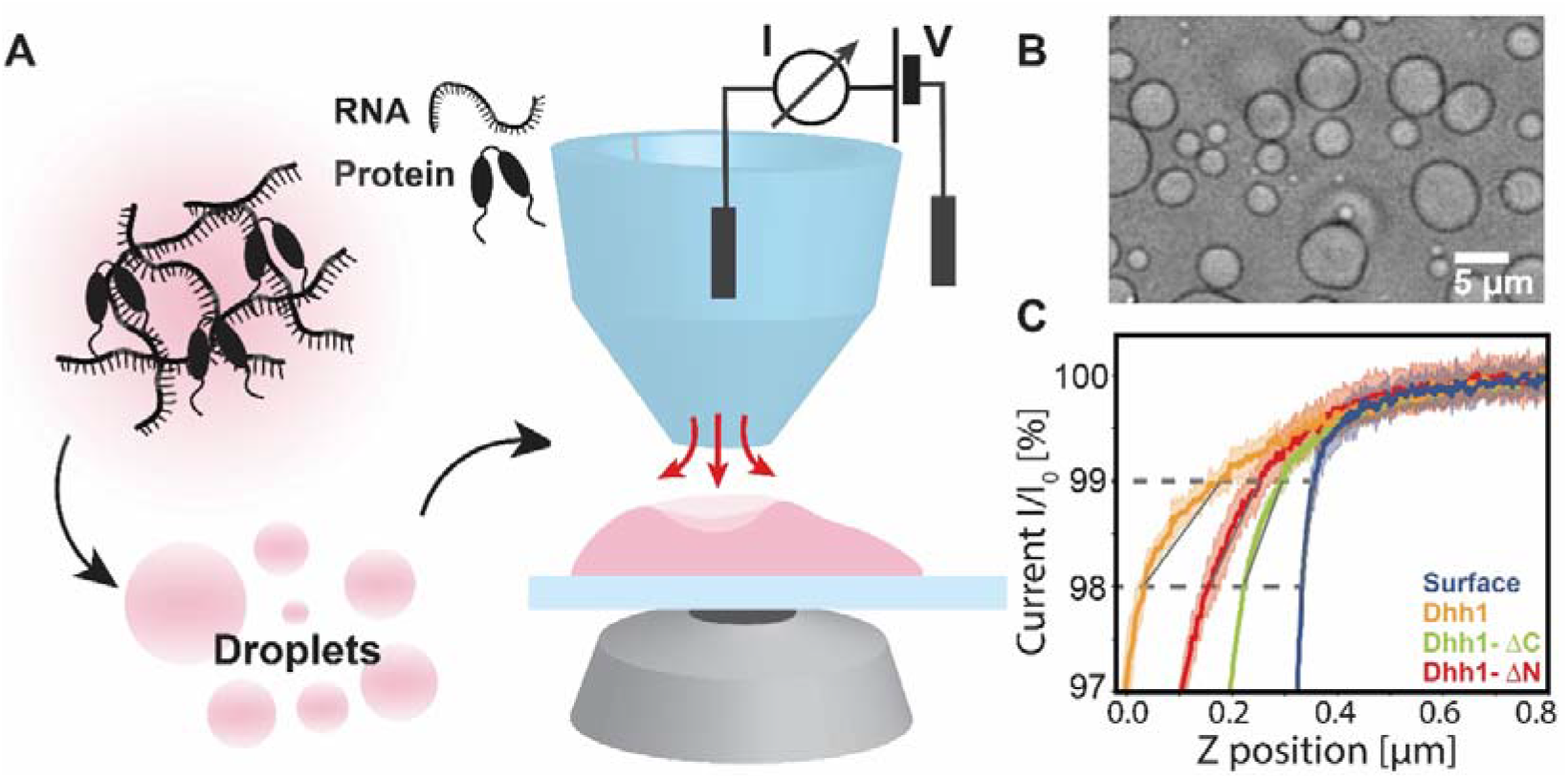
Scanning ionic conductance microscopy of biomolecular condensates. **A** Schematic of condensate assembly, in which biomolecular condensates arise through interactions between proteins and RNA and form droplets whose growth and sedimentation enable bright-field imaging, and subsequent SICM imaging allowing for 3D shape reconstruction, and local measurements of material properties. **B** Bright-field image of condensates (60× magnification) formed by Dhh1 and Poly(U) at pH 7.5, illustrating the overall sample morphology. **C** Representative SICM approach curves recorded on condensates formed by different protein variants (Dhh1, Dhh1-ΔN, and Dhh1-ΔC with Poly(U), pH 7.5), along with the corresponding surface curve highlighting their distinct mechanical responses.

To measure the stiffness of the sample, we initially generated a large topographic map of 20x20 µm with SICM topography imaging by mapping the approximate positions of the condensates on the glass surface, allowing for the alignment of the nanopipette with the condensate. After identifying a condensate to be probed, the nanopipette was then positioned directly above the center of the condensate and the trigger threshold was changed to 98% of the ionic current of the pipette corresponding to the open nanopipette current (SICM baseline current). We recorded 10-20 approach curves on the same point and averaged them (**SI Figure 1A**). An example of such an approach curve is shown in **Figure 1C**. In addition to analyzing all condensates within the scan area, identical measurements were also carried out on the glass surface. In order to calculate stiffness, we followed the protocol as outlined in Rheinlaender et al. ^19^ In this work, the stiffness (k) was shown to be directly proportional to the Young’s modulus (E), with 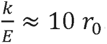. Specifically, we identified the points at which the ionic current trace reached 99% and 98% of its maximum value and measured the corresponding distance between these points (see **Figure 1C**). The slope was then defined as S = 1/d, while the Young’s modulus was calculated using:

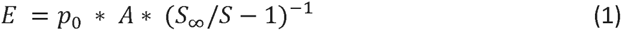

where *S*_∞_ is an approach slope measured on the glass surface, *S* is an approach slope measured on the condensate, and *p*_0_ and *A* are the pressure and shape parameters of the nanopipette, respectively. In the figures, we show the averaged relative stiffness calculated as *(S*_∞_/*S* — 1)^—1^ (**SI Figure 1B**) and then determine the Young’s modulus by multiplying it by the constant *p*_0_ *· A*, which accounts for the electroosmotic force and capillary geometry (**Figure 2A**). From here, we can calculate the stiffness by multiplying the Young’s modulus by an additional constant 10 · *r*_0_. Details of the calculation of these parameters are provided in the **Methods section**. Note that these values do not change for similar geometries and nanopipettes used, allowing the changes in the stiffness of the sample to be completely reflected in the *S*_∞_/*S* term. This permits a consistent and direct comparison between measurements on multiple samples using the same capillary another key feature of our non-contact method.

**Figure 2.**
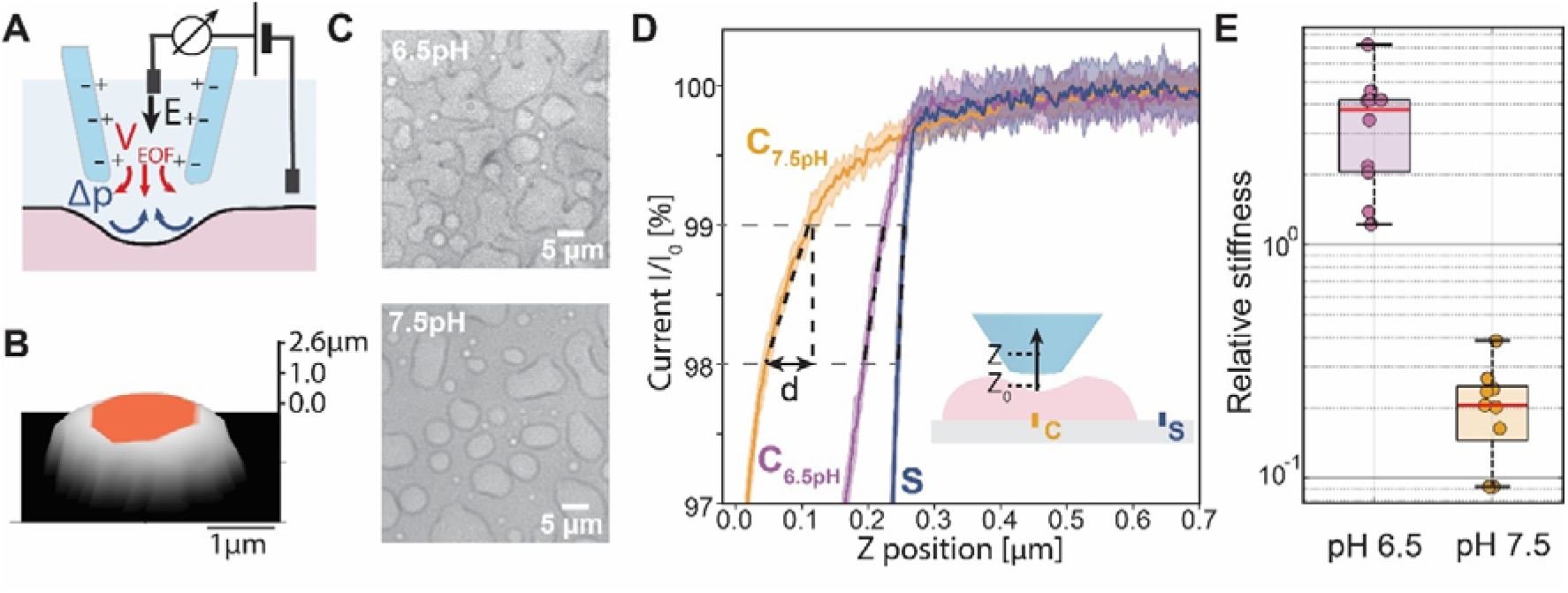
Principal of measurements of material properties - approach curves current traces with the distribution of the stiffness measurements. **A** Illustration of ionic flow conductance microscopy exerting hydrodynamic force on a soft condensate. **B** Representative 3D topography of a condensate with the orange-labeled region marking a relatively flat area suitable for stiffness measurements. **C** Bright-field image of biomolecular condensate - top at pH 6.5, bottom at pH 7.5. **D** Representative approach curves measured on condensates at pH 7.5 (orange) and pH 6.5 (magenta), as well as the surface S (blue), illustrating their different deformation responses. **E** Relative stiffness extracted for condensates at pH 6.5 and pH 7.5, revealing a pH-dependent change in condensate stiffness from (3.5 ± 0.6) to (0.20 ± 0.03) (N = 10).

Next, we mapped the influence of edges on our stiffness measurement recording the stiffness to build a cross-sectional stiffness map of the condensate. As shown in **Figure 2B** the condensates exhibit a plateau across the central regions. In contrast, closer to the edges of the condensate, the surface becomes curved and the approach angle changes, which can lead to variability in the measured stiffness (**SI Figure 2**, probing positions 0.2 μm apart). We extended our map from a cross section to an area in **Figure 2B** and highlighted the area where the obtained stiffness is consistent. We observed that when the height variation from the change in curvature is less than 8%, the measured stiffness remains unchanged (see **SI Figure 2C**). For a condensate with a height of 2 µm, this corresponds to probing approximately 0.5 µm from the edge. To allow for comparison across different conditions we collected data from this central region.

### Effect of pH on condensate stiffness

Biomolecular condensates are known to be affected by changes in pH, and specifically, Dhh1 condensation was shown to be promoted *in vitro* by lowering the pH ^17,34^. Other parameters, such as ionic strength, temperature, and buffer composition, also influence condensate formation. In this work, we focus on pH, as it leads to pronounced differences in material properties of P bodies in yeast cells and is physiologically relevant as the intracellular pH generally ranges between 6.0 and 7.5. Thus, we explored the material properties of Dhh1 condensates at different pH to probe the robustness of our methodology. The height and size distributions of the biomolecular condensates at either pH 6.5 or 7.5 were similar (**Figure 2 C**), spanning in a height range of 0.4–4 µm. However, as described above, we performed stiffness measurements on condensates with heights between 1.7 and 4 µm (see **SI Figure 3A** and **3B**) that are most frequent occurring in both conditions and to avoid edge effects. We observe a 17-fold increase in the relative stiffness at pH 6.5 (3.5 ± 0.6) compared to pH 7.5 (0.20 ± 0.03) (N=10 for both conditions), (**Figure 2 D** and **E**). For our chosen nanopipette geometry and voltage, using the calibration by Rheinlaender et. al.^21,35^, we calculated that the Young’s modulus is (1.5 ± 0.3) kPa at pH 6.5 and (0.09 ± 0.01) kPa at pH 7.5, and the stiffness is (1.4 ± 0.3) 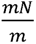 at pH 6.5 and (0.081 ± 0.009) 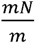 at pH 7.5.

**Figure 3.**
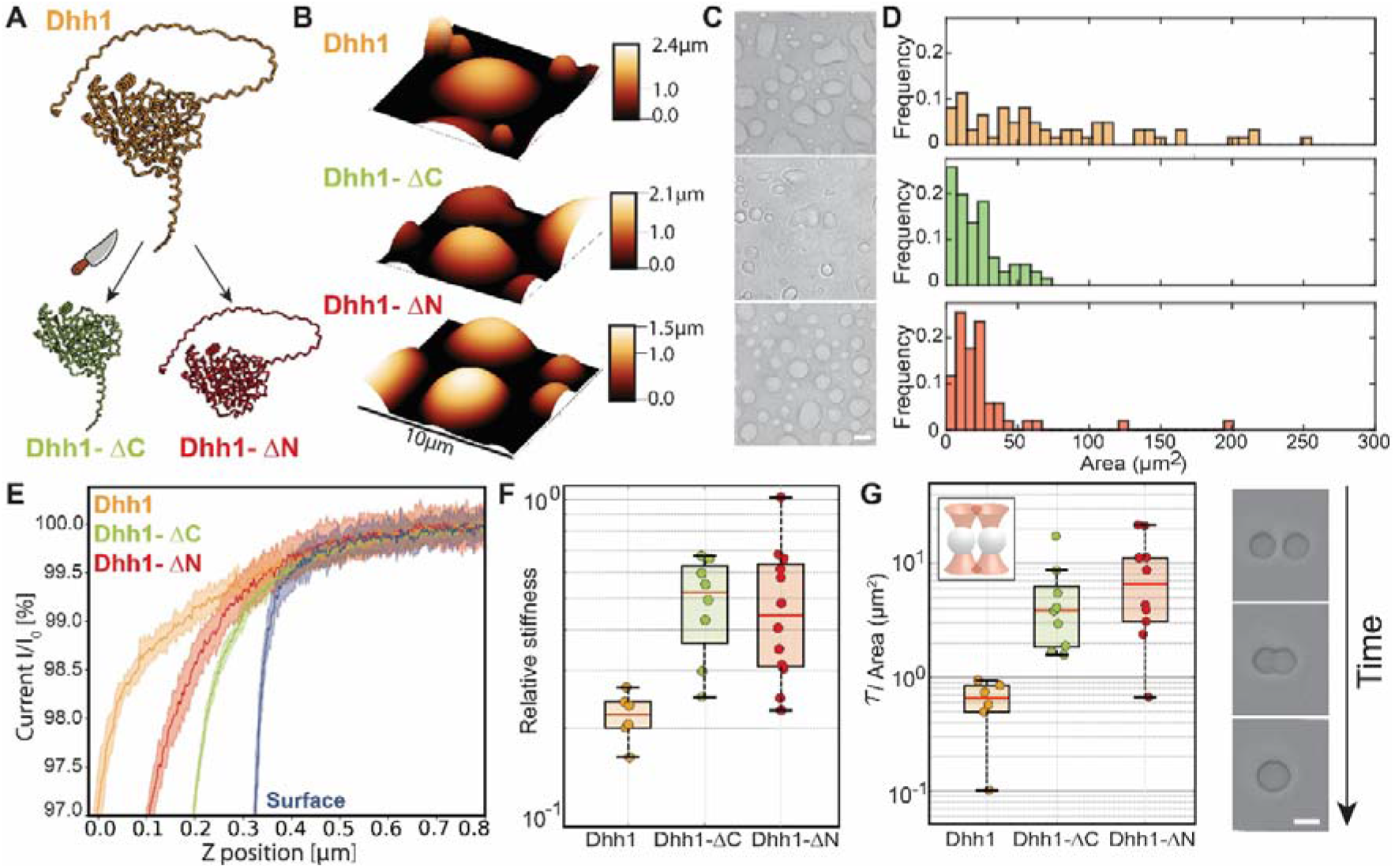
Different variants of Dhh1 protein and their condensate properties. **A** Schematic of the full length Dhh1 and the generation of its CÖterminal and NÖterminal truncation variants, Dhh1-ΔC and Dhh1-ΔN. **B** SICM 3D topography of condensates formed by Dhh1, Dhh1-ΔC and Dhh1-ΔN. **C** Bright-field images of Dhh1, Dhh1-ΔC and Dhh1-ΔN (from top to bottom, respectively) with corresponding size histogram (D), showing the distribution of projected condensate areas for each variant. Scale is 5 µm. **E** Representative approach curves recorded on the condensates made from each protein variant, highlighting their distinct mechanical responses. **F** Relative stiffness of condensates formed by the Dhh1 (N = 8), Dhh1-ΔC (N = 10) and Dhh1-ΔN (N =12). **G** Condensate fusion time measured using optical tweezers by the Dhh1 (N = 6), Dhh1-ΔC (N = 9), and Dhh1-ΔN (N = 10) proteins, averaged by condensate area and sample video stills highlighting the process. Scale bar corresponds to 5 µm.

All measurements were performed using the same nanopipette, ensuring identical capillary geometry across conditions. Because the stiffness is extracted from the relative slope *S*_∞_/*S* part of the probe-dependent contribution to the electroosmotic pressure is normalized. In addition, the pH variation explored here (6.5–7.5) is relatively small, and the buffer composition and ionic strength were kept constant, making large variations in electroosmotic mobility unlikely. Indeed, repeated measurements performed with slightly different geometries yielded similar stiffness values for condensates under the same conditions (see SI Figure 4). Therefore, the observed differences are primarily attributed to changes in condensate mechanical properties rather than variations in the electroosmotic perturbation.

**Figure 4.**
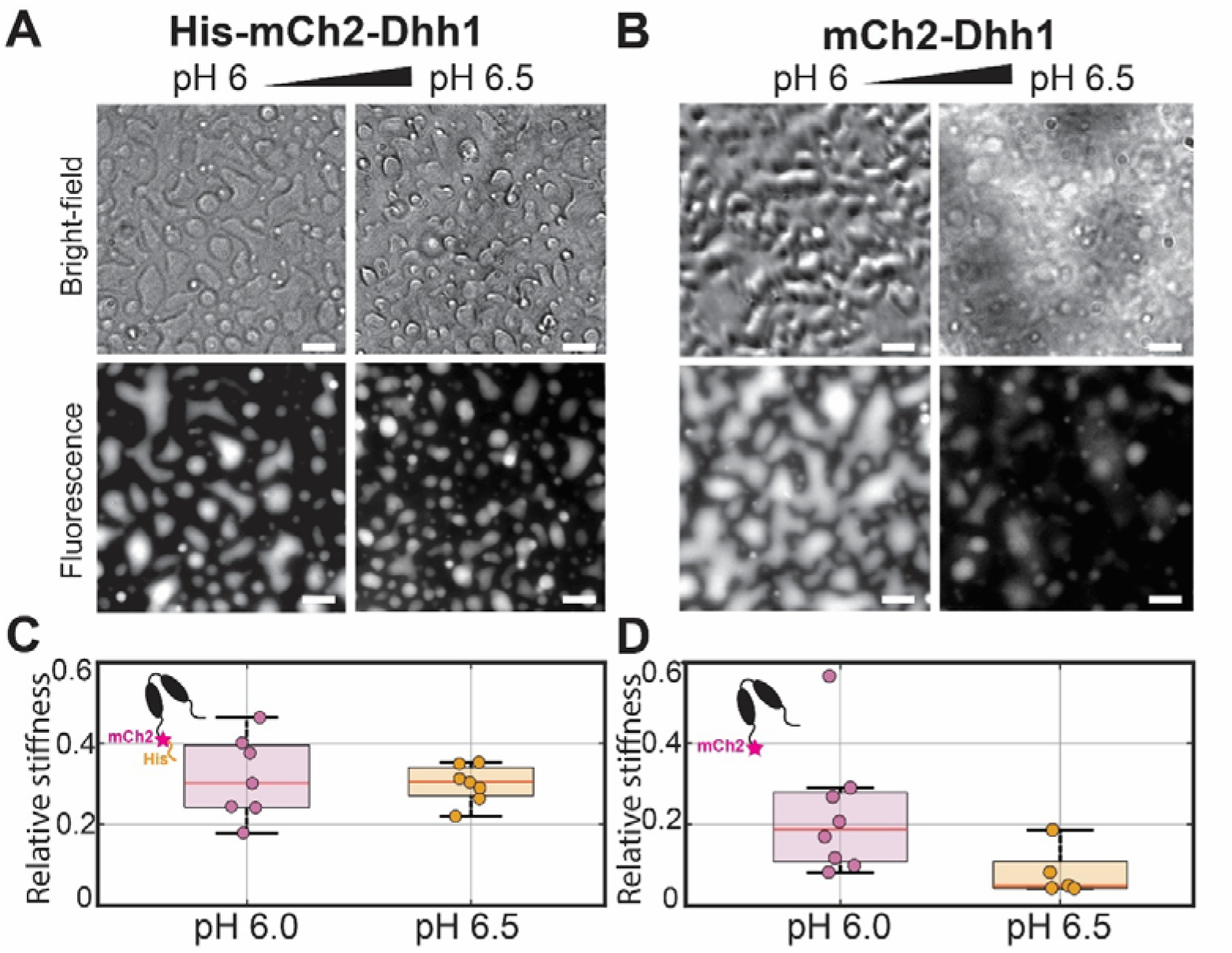
Fluorescently labeled Dhh1 protein condensates. **A** Images of His–mCh2–Dhh1 condensates at pH 6.0 (left) and pH 6.5 (right). **B** Images of mCh2–Dhh1 condensates at pH 6.0 (left) and pH 6.5 (right). Scale represents 5µm. Top: bright-field images (BF); bottom: fluorescence images (FL). **C** Relative stiffness of His-mCh2-Dhh1 condensates at pH 6.0 and pH 6.5 (N = 7). **D** Relative stiffness of mCh2-Dhh1 condensates at pH 6.0 (N = 8) and pH 6.5 (N = 6).

### Stiffness changes among structural mutants of Dhh1 (Dhh1-ΔC and Dhh1-ΔN)

To further explore our methodology, we measured the stiffness of condensates formed by Dhh1 variants that lack either N- or C-terminal disordered tails (Dhh1-ΔN, and Dhh1-ΔC), as they were previously shown to influence Dhh1 condensation (**Figure 3A**)^18^. First, we characterized the height and size of the condensates by looking at SICM topography images. The SICM topographic maps revealed that the different truncation constructs gave rise to substantially different sizes and morphology (**Figure 3B),** which was also confirmed by optical microscopy (**Figure 3C**). **Figure 3D** plots the distribution of condensate sizes of the 3 different constructs from the optical microscopy image. Whereas Dhh1-ΔC condensates exhibited a great diversity of irregular shapes, the Dhh1-ΔN condensates displayed an ellipsoid shape. Condensates formed by full-length Dhh1 exhibited a mix of both. In addition, Dhh1-ΔC condensates displayed generally smaller average sizes and height compared to Dhh1 full-length condensates. Stiffness measurements of each Dhh1 construct showed a gentler slope for Dhh1 full-length condensates compared to condensates formed by Dhh1-ΔN or Dhh1-ΔC, suggesting that the loss of either tail increases total stiffness (**Figure 3E).** We verified our data by measuring a sample population for all 3 variants (see **Figure 3F**). The resulting relative Young’s modulus values are (0.23 ± 0.02) for Dhh1 (N=8), (0.50 ± 0.05) for Dhh1-ΔC (N=10), and (0.49 ± 0.07) for Dhh1-ΔN (N=12). We calculated that the Young’s modulus is (0.090 ± 0.008) kPa, (0.20 ± 0.02) kPa, and (0.19 ± 0.02) kPa, and the stiffness is (0.081 ± 0.007) 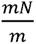, (0.18 ± 0.02) 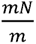, and (0.17 ± 0.02) 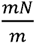, respectively for Dhh1, Dhh1-ΔC, Dhh1-ΔN.

To corroborate our SICM stiffness measurements, we performed fusion assays using optical tweezers. Fusion assays have been a standard tool employed by the field to obtain information on the material properties of various biomolecular condensates^7,8,12^. Briefly, two biomolecular condensates are trapped by the laser beam and brought closer together, and allowed to fuse and relax into a single sphere. The force curves are recorded, and the resulting curve is fitted to obtain the fusion time. This time parameter is a feature of condensate viscosity and surface tension normalized by the surface area of the condensate^4,8^. Fusion assay measurements can be compared with SICM stiffness measurements, because viscosity and stiffness are coupled in viscoelastic materials (linearly related in the Maxwell model){Citation}. We measured the fusion times of Dhh1, Dhh1-ΔN, and Dhh1-ΔC to obtain normalized fusion times a 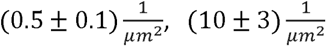, 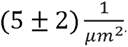, respectively (shown in **Figure 3G**). In agreement with our SICM results Dhh1 full-length condensates exhibit viscosity values that are lower than those of condensates formed by Dhh1-ΔN and Dhh1-ΔC.

### Influence of fluorescent tags on Dhh1 proteins

Previous studies have shown that fluorescent labels can significantly influence the material properties of biomolecular condensates, altering their behavior both *in vivo* and *in vitro*^17,34^. We therefore also, compared condensates formed with N-terminal tagged mCherry2-Dhh1 (mCh2–Dhh1) or doubly tagged His–mCh2–Dhh1 containing an additional array of six histidine before the mCh2 tag. Condensate formation was confirmed with both optical and fluorescence microscopy (**Figure 4A and 4B**) at two different pH conditions (pH 6.0 and pH 6.5). Previous studies by Fatti et al.^17^ using FRAP showed that at pH 6.0 and 6.5, there are differences between the recovery time and hence the diffusivity between condensates and their environment. We hypothesized that this indicates a change in the viscosity of the condensates between these two different pH levels. The images are shown in **Figure 4A** for His–mCh2–Dhh1, and **Figure 4B** for mCh2–Dhh1 taken at two different pH conditions (6.0 and 6.5).

At pH 6.0, the condensates are slightly larger than those formed at pH 6.5, and once they reach a few µm in size, they begin to lose their circular shape and start fusing, even while attached to the glass surface. Bright-field images revealed that the labeled mutants form condensates at lower pH values compared to native Dhh1, suggesting changes in material properties (see **Figure 4A** and **4B**). The relative stiffness of His–mCh2–Dhh1 is similar at pH 6.0 (0.32 ± 0.04) and pH 6.5 (0.30 ± 0.02) (**Figure 4C**), whereas the stiffness of mCh2–Dhh1 changes noticeably, decreasing from (0.23 ± 0.06) at pH 6.0 to (0.08 ± 0.02) at pH 6.5 (**Figure 4D**). The Young’s modulus for His–mCh2–Dhh1 is (0.057 ± 0.007) kPa at pH 6.0 and (0.053 ± 0.004) kPa at pH 6.5, while for mCh2–Dhh1 is (0.08 ± 0.02) kPa at pH 6.0 and (0.028 ± 0.007) kPa at pH 6.5. The stiffness values are (0.08 ± 0.01) 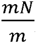, (0.074 ± 0.006) 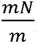, (0.12 ± 0.03) 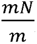, and (0.04 ± 0.01) 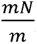, for His–mCh2–Dhh1 at pH 6.0 and 6.5, and mCh2–Dhh1 at pH 6.0 and 6.5, respectively.

Here, we observe a discrepancy of two orders of magnitude between the stiffness values measured for the tagged protein mCh2–Dhh1 and native Dhh1. The results indicate that the presence of the tag can affect the measured mechanical response, highlighting the importance of measuring unmodified systems whenever possible. Importantly, SICM enables such measurements because it does not require embedded probes, fluorescent labels, or surface attachments. As a result, the technique allows direct quantification of the mechanical properties of fully native, unperturbed condensates.

### Long-term monitoring of the stiffness of Dhh1 condensates

Since SICM is non-invasive, we explored whether our SICM methodology could be suitable to measure condensate properties over a long time period. Earlier observations by Linsenmeier et al. ^36^ showed that Dhh1 condensates undergo a stiffening when the ability to hydrolyse ATP is blocked. We therefore examined condensates formed by a mutant of Dhh1 (Dhh1^DQAD^) (**Figure 5A**) with a single amino acid mutation, which prevents ATP hydrolysis. Stiffness measurements of the condensates were performed over a period of 20 days. We noted that the topography mapping did not show significant changes in the 3D topography of the Dhh1^DQAD^ condensates at different time points, although we observed a slight increase in size between day 2 and day 20. (**Figure 5B**). For each day, after obtaining the 3D topography images, we also measured the stiffness of the condensates. On day 0, the relative stiffness was (0.23 ± 0.02), which corresponds to a Young’s modulus of (0.056 ± 0.005) kPa, and a stiffness of (0.067 ± 0.006) 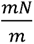. By day 1, it increased to (1.1 ± 0.1), which corresponds to Young’s modulus of (0.27 ± 0.02) kPa and stiffness of (0.32 ± 0.02) mN/m, and by day 2, it reached a plateau at (4.5 ± 0.5), corresponding to Young’s modulus of (1.1 ± 0.1) kPa and stiffness of (1.3 ± 0.1) 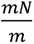. This plateau was confirmed by re-measuring the same sample on day 20, which yielded a relative stiffness of (4.0 ± 0.5), a Young’s modulus of (1.0 ± 0.1) kPa, and a stiffness of (1.2 ± 0.1) 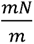, see **Figure 5C** (N = 9 for all). With this, we demonstrated that SICM is suitable for extremely long-term measurements.

**Figure 5.**
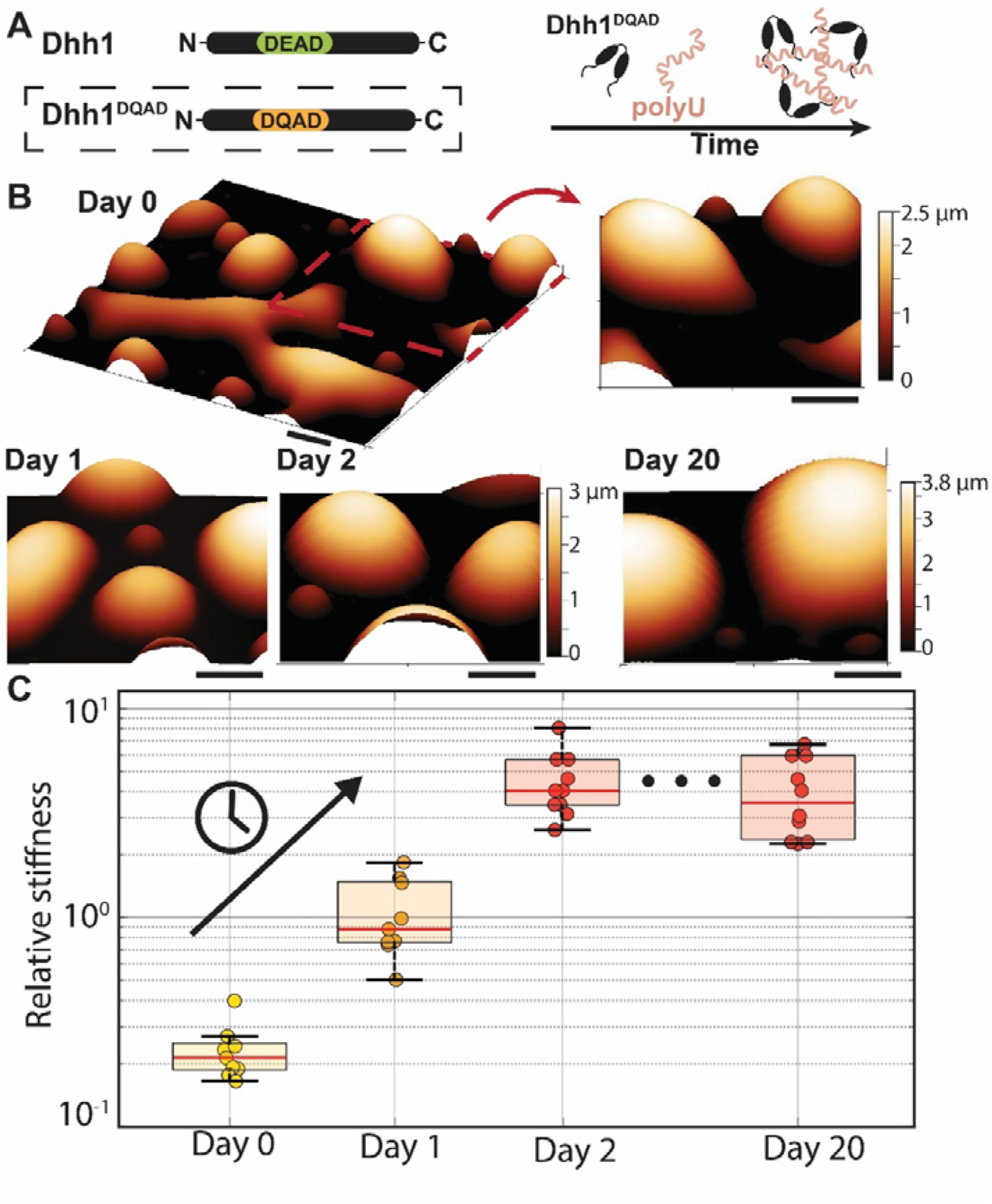
Time-dependent stiffening of the Dhh1^DQAD^ condensates. **A** Schematic of the Dhh1^DQAD^ variant in which the DQAD mutation alters ATP hydrolysis and thereby modulates interactions with RNA and condensate aging. **B** SICM 3D topography images of Dhh1^DQAD^ condensates over time. Scale is 5 µm. **C** Relative stiffness of Dhh1^DQAD^ condensates as a function of sample age, showing progressive stiffening over time (N = 10).

## Discussion

We demonstrate that SICM can be reliably used to measure the stiffness of biomolecular condensates. We strongly correlated our observations with two established techniques: optical tweezers–based fusion assays and FRAP. Our observations are in agreement with these two standard techniques, as compared in the respective **Results Section**. We note that our system offers several advantages over the standard approaches, including compatibility with physiologically relevant protein conditions, no requirement for protein modifications (i.e., no fluorescent labels), long-term measurements and monitoring of up to 20 days, and non-contact measurements that minimize sample perturbation, such as piercing, deformation, or fusion. At first glance, probing material properties with SICM is conceptually similar to viscoelastic AFM measurements, due to a similar way of scanning and probing the system, but it offers key advantages over AFM. In AFM, the sample is progressively deformed during the measurement due to interactions and sticking of the cantilever which is used as the probe which makes repeated measurements on the same condensate challenging, and the sample is progressively deformed during the measurement. By contrast, SICM offers a non-contact approach where the measurement readout arises from the electroosmotic flow that produces the indentation on the condensate from a distance, thereby eliminating contact with the condensate interface. This ensures that the nanopipettes remain robust, as demonstrated by a stable open-pore current, and allows that all measurements can be performed using the same capillary or the same batch. Using the same capillary, it is possible to directly compare the obtained stiffness result and mechanical properties across different conditions. Additionally, this robustness enables extended measurement times and allows us to measure different samples as demonstrated by our long timescale measurements up to 20 days.

However, the SICM technique as presented here also has limitations that offer future directions for development. Probing condensates smaller than ∼1 µm is challenging due to surface effects, which may alter the stiffness readout. This can be further improved through the use of smaller capillaries. In addition, similar to AFM techniques, SICM based stiffness measurements suffer from long rastering times that are needed to generate a topographic map and identify regions of interest. We tried to overcome this problem with optical microscopy to pre-position the probe. Further optimizing and automating of this process will be crucial for large-scale applications, particularly if timescales shorter than tens are to be probed. Finally, the accuracy and precision measurements of stiffness varies with the size and geometry of the used capillary, which can introduce variability when using different capillary probes. A more accurate theoretical model (accounting for the entire approach curve instead of using only two data points) or pipette shape and size standardization should be developed in future. We envision a quick calibration of the capillary size and morphology using a known standard in combination with an improved theoretical model before actual measurements on the interested analyte.

This method can be adapted and automated for a wide range of biomolecular condensates system including those with applications in controlling drug release, RNA-based therapeutics, and intracellular targeting^37,38^, where tuning condensate stiffness is important. To access stiffer material regimes and achieve higher spatial resolution, the applied nanopipette pressure can be increased and the pipette diameter reduced, thereby enhancing the resolution and range capabilities of the technique. We also envision the integration of other technical modalities demonstrated with nanopipettes, such as the use of the nanopipette for nanobiopsy^39^. This will allow an exciting combination of stiffness sensing combined with spatial transcriptomics capabilities.

## Conclusion

In summary, we characterized the material properties of Dhh1 condensates using the SICM stiffness measurement technique. First, we identified regions of the condensates and experimental conditions—specifically the approach speed and height—under which the system can be reliably applied to biomolecular condensates. These experiments were conducted under physiologically relevant protein conditions. Second, we demonstrated that the system can detect an increase in stiffness when lowering the pH from 7.5 to 6.5, mimicking physiological changes within the cell. Third, we examined how C-terminal or N-terminal truncations of the Dhh1 protein and mCh2 fluorescently labeled Dhh1 protein influence condensate formation and their material properties. Finally, we demonstrated long-term tracking of stiffness measurements up to 20 days. We corroborate our results with established techniques such as optical tweezer fusion assay measurements and FRAP. Our approach can be extended to study other biomolecular condensates and similar soft matter. We believe that this technology will further advance our understanding of the regulation of function and the physical properties of biomolecular condensates.

## Methods

### Sample preparation

#### Expression and Purification of Dhh1

Dhh1FL (pKW5049), Dhh1(32–506) (pKW5077), Dhh1(1–425) (pKW5076), Dhh1DQAD (pKW5081), and mCherry2–Dhh1FL (pKW5066) were expressed in E. coli Rosetta (DE3) (Novagen) as 6×His–TEV-tagged proteins using auto-induction medium. Cultures were grown at 37 °C in 1 L ZY complete medium to an OD600 of 0.7, after which the temperature was reduced to 20 °C and incubation continued for 20 h at 220 rpm. Cells were harvested, washed with cold PBS, flash-frozen in liquid nitrogen, and stored at −20 °C until further use. Cell pellets were resuspended in 10 mL/g of lysis buffer (20 mM HEPES-KOH, pH 7.7, 500 mM KCl, 5 mM MgCl2, 0.2% IGEPAL CA-630, 10 mM imidazole, and 2 mM β-mercaptoethanol) supplemented with 0.5 mg/mL lysozyme and 0.01 mg/mL DNase I (PanReac AppliChem), and lysed using a high-pressure homogenizer (Emulsiflex C5, Avestin). The lysate was clarified by centrifugation at 20,000 × g (SS-34 fixed-angle rotor, Sorvall), and the supernatant was filtered through a 0.45 µm PES membrane (Sarstedt) before incubation with Ni-NTA resin (Qiagen). The resin was sequentially washed with five column volumes of detergent buffer (20 mM HEPES-KOH, pH 7.7, 500 mM KCl, 5 mM MgCl2, 0.2% IGEPAL CA-630, 10 mM imidazole, 2 mM β-mercaptoethanol, 5% glycerol), high-salt buffer (20 mM HEPES-KOH, pH 7.7, 1.5 M KCl, 5 mM MgCl2, 10 mM imidazole, 2 mM β-mercaptoethanol, 5% glycerol), and imidazole buffer (20 mM HEPES-KOH, pH 7.7, 1.5 M KCl, 5 mM MgCl2, 20 mM imidazole, 2 mM β-mercaptoethanol, 5% glycerol). Proteins were eluted with 2.5 column volumes of elution buffer (20 mM HEPES-KOH, pH 7.7, 500 mM KCl, 5 mM MgCl2, 330 mM imidazole, 2 mM β-mercaptoethanol, 5% glycerol). Eluted proteins were buffer-exchanged into imidazole buffer using a PD-10 column (GE Healthcare) and incubated overnight at 10 °C with 6×His–TEV protease. The sample was then reapplied to Ni-NTA resin to remove uncleaved protein and the protease. The flow-through was concentrated using centrifugal filter units (Millipore) and further purified by size-exclusion chromatography on a Superdex 200 16/600 column (GE Healthcare) in SEC buffer (20 mM HEPES-KOH, pH 7.7, 500 mM KCl, 5 mM MgCl2, 1 mM DTT, 10% glycerol) using an ÄKTA pure system (GE Healthcare). Fractions containing the target protein were pooled, concentrated, and purity was assessed by SDS-PAGE followed by Coomassie staining (InstantBlue®, Abcam). For 6×His–mCherry2–Dhh1, the TEV cleavage step was omitted, and the protein was directly subjected to size-exclusion chromatography. Final samples were aliquoted, flash-frozen, and stored at −80 °C.

#### Biomolecular condensate and buffer preparation

A working buffer containing 100 mM KCl, 1mM MgCl_2_, 30 mM HEPES was prepared and the pH was adjusted with 5M KOH till it reached pH 7.5. The buffer was then purified with a 0.22 µm filter. For experiments at pH 6.0 or 6.5, a similar working buffer was used, except that 30 mM MES was used instead of HEPES. All buffers were stored at room temperature. If not stated explicitly, the experiments in the paper were all performed at pH 7.5.

For condensate formation, 200 µM Uracil homopolymer RNA (Microsynth) was diluted into 80 µl of working buffer. DTT was used to supplement the buffer at a final concentration of 0.5 mM. The proteins were added right before imaging to a final concentration of 5 µM. All experiments were performed at room temperature.

Biomolecular condensates undergo time-dependent ageing; it is important that imaging, SICM 3D scanning, and mechanical property measurements are performed at comparable stages of aging. When comparing the material properties of condensates of different Dhh1 mutants and at different pH, condensates were imaged within a time window of 3-5 hours after incubation, when the condensates were settled onto the glass surface. During this period, samples were handled with care to prevent evaporation and associated changes in salt concentration.

### Probe preparation

#### Nanopipette preparation

Borosilicate glass pipette (Science product, GB120F-10, length 100 mm, outside diameter 1.2 mm, and inside diameter 0.69 mm) were pulled using a CO2 laser puller (P-2000, Sutter Instruments) with the following protocol: Heat = 440, Fil = 3, Vel = 35, Del = 220, Pul = 0; Heat = 470, Fil = 4, Vel = 45, Del= 175, Pul = 250. Nanopipettes were then placed inside a plasma cleaner and filled from the back side with the needle and syringe with the chosen buffer solution. Nanopipettes fabricated in this way have a radius ranging from 90 to 140 nm.

### Experimental setup

#### SICM topography imaging

SICM is a non-contact topography imaging tool that allows for measurements in environmental conditions. To approach the surface, we used the characteristic decrease in ionic current that occurs when the nanopipette comes into proximity to the surface. During imaging, the system is triggered to record the height of the object at the position where the ionic current drops to 99% of the value. These height measurements are then combined to reconstruct a topographical image.

The protocol of the experiment is that the nanopipette is inserted in the buffer and lowered toward the surface with coordinated movement in the z-direction using both the stepper motor and the piezo stage. During the approach, a trigger is activated to stop the motion in case the measured ionic current decreases to 99% of its initial value. The approach starts with the piezo stage moving through its full range (for our system – 20 μm; with a speed of 50 μm/s). If the threshold is not reached, the piezo retracts to its minimum position, and a step motor makes a step that is 60 % of the piezo’s full range. Alternating these two setups allows us to progressively assess proximity to the surface using the piezo stage, followed by stepwise advancement with the step motor. Once the threshold is reached, the nanopipette is in the approached position (with reference to the surface) and stops moving. After capillary is approached the images is taken with hopping rate of 40-80 Hz, and a retracting height of 7 µm, data acquired at a rate of 1 MHz.

#### SICM stiffness measurements

SICM can be used to measure stiffness by applying a pressure-induced flow to the sample and analyzing its mechanical response. The applied pressure can be increased by bringing the probe closer to the sample by lowering the triggering point to 96% of the ionic current (to obtain a clear average value measured at 98%), and with the same hopping rate and retracting height. The current is then recorded and analysed.

##### Data analysis

After recording the approach curves, the data were analyzed by identifying the position at which the defined current threshold was reached and saving 0.06-0.12s (3000-6000 points) before this point (see **SI Figure 1A**). For each measurement, we averaged over 10 recorded curves by calculating the root mean square (RMS) and standard deviation. At this point, we look at the slope coefficient between 99% and 98% of the relative current *I*/*I*_0_ (see **Figure 1B**). In each experiment, data were collected from representative condensates (size of the condensate usually 1.5-4 µm). For each condensate slope coefficient, S, was extracted and compared to the surface slope coefficient *S*_∞_. The relative stiffness was then calculated and is presented as a box plot with the mean value indicated. From her Young’s modulus was subsequently obtained by multiplying it by the shape parameter A, which is 0.33 for the capillary with 2° angle^21^, and the electro-osmotic pressure calculated as 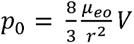, where η is the viscosity, *r* is the capillary radius, and *V* is the applied voltage^21,35^. The stiffness was calculated by multiplying it with 10 · *r*.

#### Fusion Assays

Fusion assays were performed using the NanoTracker™ 2 Optical Tweezer system (Bruker). The same method mentioned in the Biomolecular condensate and buffer preparation section was used for condensate formation on a #1.5 round glass coverslip (30 mm in diameter), with a custom-cut (square with a circular cut-out) double-sided adhesive silicone spacer (0.36 mm in thickness). Once the condensate solution was added into the center of the cut-out, an identical coverslip was placed over it to create a sealed sample chamber. For all measurements, stiffness values of ∼0.06⍰pN/nm were used. Coalescence was initiated by moving one trapped condensate closer to another, until a characteristic spike in normalized force was observed. Both force-time traces and .avi video files were recorded for each event, with the FOV shifting between measurements to dampen significant temperature changes. Each dataset lasted 15 to 45 minutes.

##### Data analysis

The τ values were determined by fitting the relaxation force data to a stretched exponential function *F*(*t*) = *a + (b − a)[1 − e^−^*^((*t−t*_0_)/(^*^τ^*^)^*^B^*] on MATLAB. The diameter of the droplets was determined from the .avi files using a custom MATLAB script to normalize τ by area.

#### Fluorescence and bright field imaging

To assess the size and shape of the condensate, images were acquired using a 60× objective in both bright-field and fluorescence modes. Fluorescence imaging was performed using a mercury vapor lamp (X-Cite 120Q) equipped with an excitation filter of 513 - 556 nm and an emission filter of 570 - 613 nm. For confocal imaging, we used a Leica TCS SP8 with an objective (×63 magnification, 1.4 NA).

##### Size measurements

The two-dimensional projected area of the condensates was measured using ImageJ. Condensate boundaries were identified by edge detection, and the area was calculated based on a pixel size of 135 nm × 135 nm. Pixel size calibration was performed using 1 μm reference beads.

## Contributions

W.Y., A.R., and H.M. oversaw the project and designed the experiments. H.M. prepared samples and performed SICM experiments, data analysis, and created the figures. E.F. and K.W. purified the protein and provided expertise in protein handling. K.P. and W.Y. performed the optical tweezers fusion experiments and analyzed the data. M.P., Z.A., J.S., and H.M. contributed to the optimization of the SICM software and hardware. H.M. and W.Y. wrote the manuscript, with input from A.R. and K.W., and all coauthors. All authors contributed to discussions of the results and provided feedback on the manuscript.

## Acknowledgement

H.M, E.F. K.W., W..Y. and A.R would like to thank funding from SNSF Grant CRSII5_193740. H.M. and A.R. acknowledges support from the European Research Council under grant no. 101020445—2D-LIQUID. We would like to acknowledge Prof. Georg Fantner for valuable advice on best practices for SICM use, access to the instrument and for providing insightful comments on the manuscript.

## Notes

### Competing Interest Statement

The authors have declared no competing interest.

### Summary of Updates

We corrected a minor typo in the manuscript.

## References

1. Shin, Y. & Brangwynne, C. P. Liquid phase condensation in cell physiology and disease. Science 357, eaaf4382 (2017).

2. Banani, S. F., Lee, H. O., Hyman, A. A. & Rosen, M. K. Biomolecular condensates: organizers of cellular biochemistry. Nat Rev Mol Cell Biol 18, 285–298 (2017).

3. Ross, C. A. & Poirier, M. A. Protein aggregation and neurodegenerative disease. Nat Med 10, S10–S17 (2004).

4. Patel, A. et al. A Liquid-to-Solid Phase Transition of the ALS Protein FUS Accelerated by Disease Mutation. Cell 162, 1066–1077 (2015).

5. Ray, S. et al. α-Synuclein aggregation nucleates through liquid–liquid phase separation. Nat. Chem. 12, 705–716 (2020).

6. Visser, B. S., Lipiński, W. P. & Spruijt, E. The role of biomolecular condensates in protein aggregation. Nat Rev Chem 8, 686–700 (2024).

7. Ibrahim, K. A., Naidu, A. S., Miljkovic, H., Radenovic, A. & Yang, W. Label-Free Techniques for Probing Biomolecular Condensates. ACS Nano 18, 10738–10757 (2024).

8. Michieletto, D. & Marenda, M. Rheology and Viscoelasticity of Proteins and Nucleic Acids Condensates. JACS Au 2, 1506–1521 (2022).

9. Jawerth, L. M. et al. Salt-Dependent Rheology and Surface Tension of Protein Condensates Using Optical Traps. Phys. Rev. Lett. 121, 258101 (2018).

10. Li, X., van der Gucht, J., Erni, P. & de Vries, R. Active microrheology of protein condensates using colloidal probe-AFM. Journal of Colloid and Interface Science 632, 357–366 (2023).

11. Naghilou, A., Armbruster, O. & Mashaghi, A. Scanning probe microscopy elucidates gelation and rejuvenation of biomolecular condensates. CR-PHYS-SC 6, (2025).

12. Jawerth, L. et al. Protein condensates as aging Maxwell fluids. Science 370, 1317–1323 (2020).

13. Alshareedah, I. et al. Interplay between Short-Range Attraction and Long-Range Repulsion Control s Reentrant Liquid Condensation of Ribonucleoprotein–RNA Complexes. J. Am. Chem. Soc. 141, 14593–14602 (2019).

14. Català-Castro, F. et al. Measuring age-dependent viscoelasticity of organelles, cells and organisms with time-shared optical tweezer microrheology. Nat. Nanotechnol. 20, 411–420 (2025).

15. Taylor, N. O., Wei, M.-T., Stone, H. A. & Brangwynne, C. P. Quantifying Dynamics in Phase-Separated Condensates Using Fluorescence Recovery after Photobleaching. Biophysical Journal 117, 1285–1300 (2019).

16. Lorén, N. et al. Fluorescence recovery after photobleaching in material and life sciences: putting theory into practice. Quarterly Reviews of Biophysics 48, 323–387 (2015).

17. Fatti, E., Khawaja, S. & Weis, K. The dark side of fluorescent protein tagging – the impact of protein tags on biomolecular condensation. 2024.11.23.624970 Preprint at 10.1101/2024.11.23.624970 (2024).

18. Hondele, M. et al. DEAD-box ATPases are global regulators of phase-separated organelles. Nature 573, 144–148 (2019).

19. Rheinlaender, J. & E. Schäffer, T. Mapping the mechanical stiffness of live cells with the scanning ion conductance microscope. Soft Matter 9, 3230–3236 (2013).

20. Rheinlaender, J. & Schäffer, T. E. Measuring the Shape, Stiffness, and Interface Tension of Droplets with the Scanning Ion Conductance Microscope. ACS Nano 18, 16257–16264 (2024).

21. Rheinlaender, J. & Schäffer, T. E. Quantifying and Utilizing Electroosmotic Flow for Mechanical Measurements with the Scanning Ion Conductance Microscope. Anal. Chem. 97, 22541–22547 (2025).

22. Hansma, P. K., Drake, B., Marti, O., Gould, S. A. C. & Prater, C. B. The Scanning Ion-Conductance Microscope. Science 243, 641–643 (1989).

23. Leitao, S. M. et al. Time-Resolved Scanning Ion Conductance Microscopy for Three-Dimensional Tracking of Nanoscale Cell Surface Dynamics. ACS Nano 15, 17613–17622 (2021).

24. Sánchez, D. et al. Localized and non-contact mechanical stimulation of dorsal root ganglion sensory neurons using scanning ion conductance microscopy. Journal of Neuroscience Methods 159, 26–34 (2007).

25. Scanning Ion Conductance Microscopy. vol. 3 (Springer International Publishing, Cham, 2022).

26. Simeonov, S. & Schäffer, T. E. High-speed scanning ion conductance microscopy for sub-second topography imaging of live cells. Nanoscale 11, 8579–8587 (2019).

27. Korchev, Y. E. et al. Specialized scanning ion-conductance microscope for imaging of living cells. Journal of Microscopy 188, 17–23 (1997).

28. Rheinlaender, J. & Schäffer, T. E. Image formation, resolution, and height measurement in scanning ion conductance microscopy. J. Appl. Phys. 105, 094905 (2009).

29. Kolmogorov, V. S. et al. Mapping mechanical properties of living cells at nanoscale using intrinsic nanopipette–sample force interactions. Nanoscale 13, 6558–6568 (2021).

30. Linder, P. & Jankowsky, E. From unwinding to clamping - the DEAD box RNA helicase family. Nat Rev Mol Cell Biol 12, 505–516 (2011).

31. Parker, R. & Sheth, U. P Bodies and the Control of mRNA Translation and Degradation. Molecular Cell 25, 635–646 (2007).

32. Chen, C.-C., Zhou, Y. & Baker, L. A. Scanning Ion Conductance Microscopy. Annual Review of Analytical Chemistry 5, 207–228 (2012).

33. Rheinlaender, J. & Schäffer, T. E. An Accurate Model for the Ion Current–Distance Behavior in Scanning Ion Conductance Microscopy Allows for Calibration of Pipet Tip Geometry and Tip–Sample Distance. Anal. Chem. 89, 11875–11880 (2017).

34. Dörner, K. et al. Fluorescent protein and peptide tags alter condensate formation and dynamics in vivo and in vitro. EMBO Rep 27, 89–121 (2026).

35. Kestin, J., Khalifa, H. E. & Correia, R. J. Tables of the dynamic and kinematic viscosity of aqueous KCl solutions in the temperature range 25–150 °C and the pressure range 0.1–35 MPa. Journal of Physical and Chemical Reference Data 10, 57–70 (1981).

36. Linsenmeier, M. et al. Dynamic arrest and aging of biomolecular condensates are modulated by low-complexity domains, RNA and biochemical activity. Nat Commun 13, 3030 (2022).

37. Han, T. W., Portz, B., Young, R. A., Boija, A. & Klein, I. A. RNA and condensates: Disease implications and therapeutic opportunities. Cell Chemical Biology 31, 1593–1609 (2024).

38. Xing, Z. et al. Intracellular mRNA phase separation induced by cationic polymers for tumor immunotherapy. J Nanobiotechnol 20, 442 (2022).

39. Sahota, A. et al. Spatial and Temporal Single-Cell Profiling of RNA Compartmentalization in Neurons with Nanotweezers. ACS Nano 19, 18522–18533 (2025).

